# Early candidate urine biomarkers for detecting Alzheimer’s disease before beta amyloid plaque deposition in an APP (swe)/PSEN1^dE9^ transgenic mouse model

**DOI:** 10.1101/258921

**Authors:** Fanshuang Zhang, Jing Wei, Xundou Li, Chao Ma, Youhe Gao

**Affiliations:** Department of Pathophysiology, Institute of Basic Medical Sciences Chinese Academy of Medical Sciences, School of Basic Medicine Peking Union Medical College, Beijing, 100005, China; Department of Pathology, National Cancer Center/Cancer Hospital, Chinese Academy of Medical Sciences and Peking Union Medical College, Beijing, 100021, China; Department of Biochemistry and Molecular Biology, Beijing Normal University, Gene Engineering Drug and Biotechnology Beijing Key Laboratory, Beijing, 100875, China; Institute of Basic Medical Sciences, Chinese Academy of Medical Sciences, Department of Human Anatomy, Histology and Embryology, Neuroscience Center; Joint Laboratory of Anesthesia and Pain, School of Basic Medicine, Peking Union Medical College, Beijing, 100005, China

**Keywords:** Alzheimer’s disease (AD), urine proteome, early diagnosis, APP (swe)/PSEN1^dE9^

## Abstract

Alzheimer’s disease (AD) is an incurable age-associated neurodegenerative disorder that is characterized by irreversible progressive cognitive deficits and extensive brain damage. The identification of candidate biomarkers before beta amyloid plaque deposition occurs is therefore of great importance for the early intervention of AD. Urine, which is not regulated by homeostatic mechanisms, theoretically accumulates changes associated with AD earlier than cerebrospinal fluid and blood. In this study, an APP (swe)/PSEN1dE9 transgenic mouse model was used to identify candidate biomarkers for early AD. Urine samples were collected from 4-, 6-, and 8-month-old transgenic mouse models, and the urinary proteomes were profiled using liquid chromatography coupled with tandem mass spectrometry (LC-MS/MS). The levels of 33 proteins differed significantly between wild-type and 4-month-old mice, which had not started to deposit beta amyloid plaque. Among these proteins, 16 have been associated with the mechanisms of AD, while 9 have been suggested as AD biomarkers. Our results indicated that urine proteins enable detecting AD before beta amyloid plaque deposition, which may present an opportunity for intervention.

## Introduction

Alzheimer’s disease (AD) is a chronic age-associated neurodegenerative disorder associated with cognitive impairment and progressive dementia[1]. As the pathological process leading to AD begins decades before clinical symptoms, finding early clues in the early stages of AD, especially before beta amyloid plaque deposition, is urgent in today’s AD research. Most candidate AD biomarkers came from cerebrospinal fluid (CSF) and blood studies; for example, tau levels increase in the CSF of most AD patients, and stable miRNAs in human serum are potentially valuable novel biomarkers for the diagnosis of AD. However, beta amyloid deposition can appear in AD patients, even in the stage 1 of preclinical AD, and can be detected by positron emission tomography (PET) amyloid imaging or CSF amyloid beta levels[1].

Biomarkers are measurable changes associated with disease. Unlike blood, which is stable because of homeostasis mechanisms, urine can accumulate many kinds of changes; some of these changes are associated with disease and will become biomarkers[2,3]. In addition, urine is a resource for biomarker discovery that easily and non-invasively collected from AD patients. As a sensitive biomarker source, urine therefore might reflect pathological changes, especially in the early stages of disease[4]. However, whether time-course analyses of urine proteins at different AD phases can reveal sensitive biomarker clues before beta amyloid deposition is unclear, as urinary proteins are easily affected by complicated factors such as medicine and diet, especially in human samples. Using animal models is therefore the most efficient way to establish an association with disease, as animal models can reduce the influence of genetic and environmental factors on the urine proteome to the greatest degree[2]. In addition, animal models can represent the early stages of disease without clinical symptoms as it is impossible to clinically collect such samples[5].

Several studies have applied proteomic[6] and metabolomic[7,8] analyses to characterize candidate biomarkers in mouse models. However, these studies used either plasma or brain tissues, and analyses of these materials were not sensitive enough for the early diagnosis of AD (before beta amyloid plaque deposition). Other studies have illustrated urine markers using samples from AD patients[9]. However, all these studies were conducted after beta amyloid deposition had appeared in brain tissue and cognitive impairment or movement disorder symptoms had occurred. CSF has a greater homeostatic priority than blood and changes less than blood. Urine may thus be a better choice for the detection of earlier biomarkers, even for brain diseases[10]. In addition, the limited studies about urine-based biomarkers of brain diseases suggest that urine is an ideal potential source for biomarkers in brain diseases[11]. For example, urine biomarkers in an astrocytoma rat model can reflect early astrocytoma changes before Magnetic Resonance Imaging (MRI) and thus gives potential clues for the early clinical diagnosis of astrocytoma patients[12].

Amyloid precursor protein (APP) (swe)/PSEN1^dE9^ transgenic mice, overexpressing mutant APP and PS1 (APP/PS1), have been widely used as a model of AD to elucidate the pathogenic processes of the disease and to investigate candidate biomarkers[13]. Deposition of beta amyloid in the hippocampus of 6-month-old mice occasionally occurred[14,15], and a mass of beta amyloid had been deposited in the hippocampus of 8-month-old mice [16,17]. Compared with wild-type mice, the 8-month-old transgenic mice were mainly impaired in the foreground fear conditioning test and in plasticity to acquire strategies to swim toward a cued escape platform[18]. The 12-month-old transgenic mice showed spatial learning deficits as well as long-term contextual memory deficits and large deposits of aggregated insoluble beta amyloid[19]. The cognitive abnormality of APP (swe)/PSEN1^dE9^ transgenic mice appeared at 4 months, when beta amyloid plaques had not yet deposited[20]. Potential candidate urinary biomarkers that appear before brain pathology can thus be identified by using 4-month-old APP (swe)/PSEN1^dE9^ transgenic mice. Figure 1 presents the characteristics of behavioral and pathological profiles of different stages of the APP (swe)/PSEN1^dE9^ transgenic mouse model.

**Figure 1.**
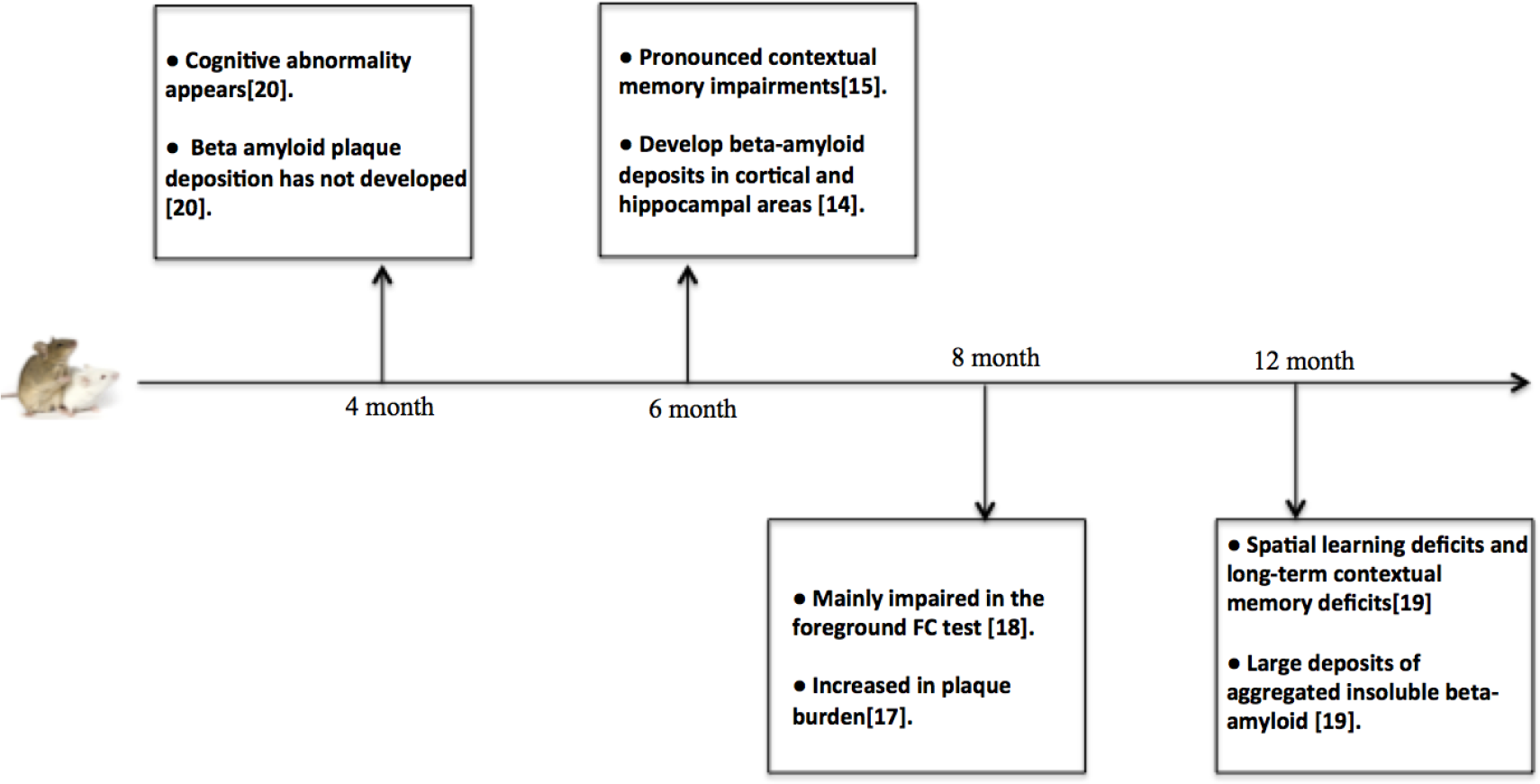
Characteristics of behavioral and pathological profiles of different stages of APP (swe)/PSEN1^dE9^ transgenic mice

In this study, the APP (swe)/PSEN1^dE9^ transgenic mouse model was used to identify candidate early AD biomarkers. To identify changes in the urinary proteome during AD development, we collected and analyzed urine samples of APP (swe)/PSEN1^dE9^ transgenic mice at three growth stages (4-, 6- and 8-month-old). The workflow of the proteomic analysis in this study is shown in Figure 2. By using label-free and TMT-labeling proteomics analysis, we analyzed differential urine biomarkers from different stages of AD.

**Figure 2.**
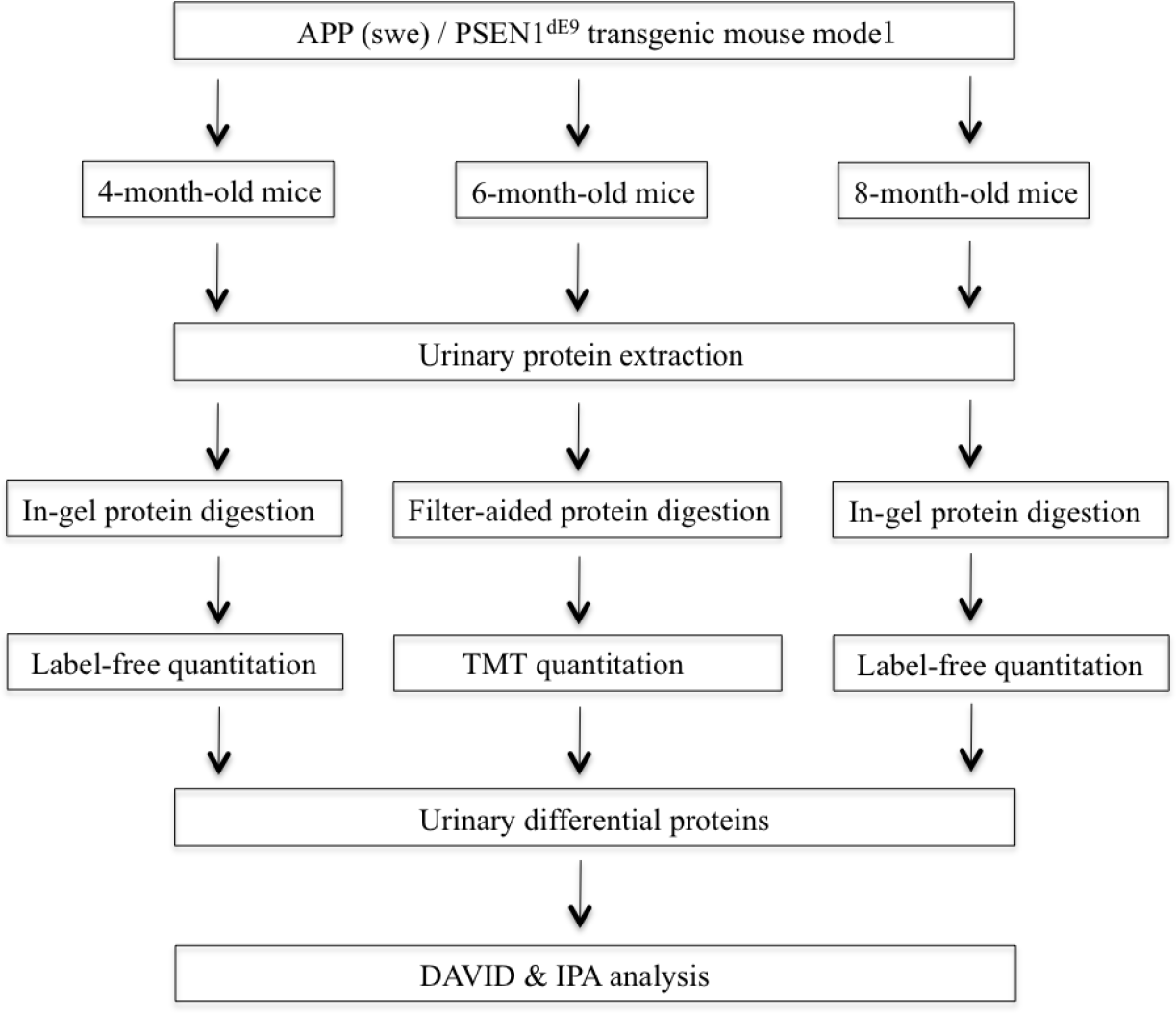
Workflow of urinary proteomics in APP (swe)/PSEN1^dE9^ transgenic mouse model. Urine samples were collected from the control and transgenic groups. The proteins were analyzed using liquid chromatography coupled with tandem mass spectrometry (LC-MS/MS) identification. Differential proteins were analyzed by DAVID and IPA.

**Figure 3.**
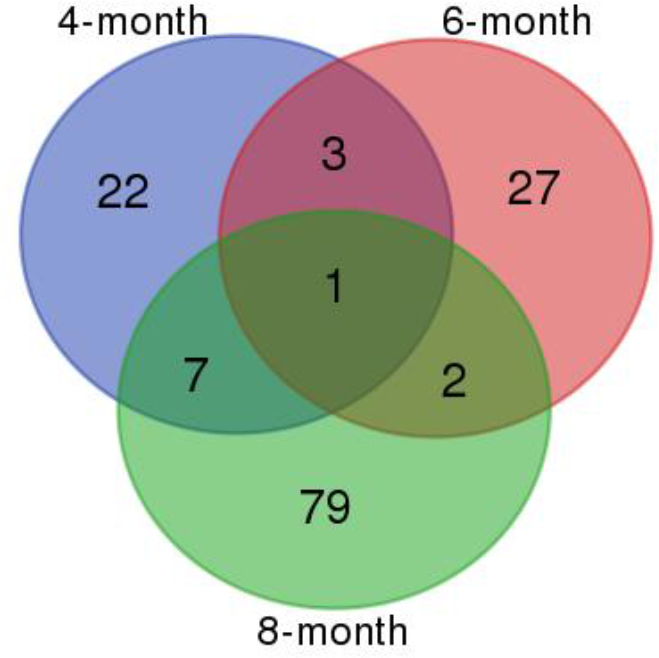
Urinary proteins with significantly different levels in the 4-, 6- and 8-month-old mice and wild-type mice

## Materials and methods

### Experimental animals

APP (swe)/PSEN1^dE9^ transgenic mice and wild-type mice were used in this study. All animal protocols governing the experiments in this study were approved by the Institute of Basic Medical Sciences Animal Ethics Committee, Peking Union Medical College (Approved ID: ACUC-A02-2014-008). All animals were maintained with a standard laboratory diet at a controlled indoor temperature (22 ± 1°C) and humidity (65 ~ 70%) and with a 12-h light-dark cycle. The study was performed according to the guidelines developed by the Institutional Animal Care and Use Committee of Peking Union Medical College. All efforts were made to minimize suffering.

### Urinary protein extraction

Urine samples from 4-month-, 6-month- and 8-month-old mice in transgenic and wide-type control groups were collected in metabolic cages. During urine collection, all rats were given free access to water without food to avoid contamination.

Urine was centrifuged at 2,000×g for 15 min at 4°C. After the cell debris had been removed, the supernatant was centrifuged at 12,000×g for 15 min at 4°C. Three volumes of acetone were added after the pellets had been removed, and precipitation was allowed to occur at 4°C. After the supernatant was removed, the pellets were resuspended in lysis buffer (8 M urea, 2 M thiourea, 25 mM dithiothreitol (DTT) and 50 mM Tris). Protein concentrations were measured using the Bradford method.

### In-gel protein digestion

Proteins were digested with trypsin (Trypsin Gold, Mass Spec Grade, Promega, Fitchburg, Wisconsin, USA) using in-gel protein digestion[21]. Protein samples were separated by SDS-PAGE. The gel was rinsed three times, 5 minutes each, in pure water and stained for 30 minutes in Coomassie brilliant blue at room temperature, after which the staining solution was discarded. The gel was destained for 1 hour in pure water at room temperature. When destaining was complete, the solution was discarded. A clean razor blade was used to cut the protein bands from the gel. The gel slices were then placed into a 0.5-ml microcentrifuge tube that had been prewashed twice with 50% acetonitrile (ACN)/0.1% trifluoroacetic acid (TFA). The gel slices were destained twice with 0.2 ml of 100 mM _NH4HCO3_/50% ACN for 45 minutes each at 37°C to remove the Coomassie brilliant blue stain. The gel slices were dehydrated for 5 minutes at room temperature in 100 μl of 100% ACN. At this point, the gel slices were much smaller than their original size and whitish or opaque in appearance. The gel slices were dried in a Speed Vac^®^ for 10 minutes at room temperature to remove the ACN. Trypsin Gold was resuspended at 1 µg/µl in 50 mM acetic acid and then diluted in 40 mM NH_4_HC0_3_/10% ACN to 20 µg/ml. The gel slices were preincubated in a minimal volume (10–20 µl) of the trypsin solution at room temperature (do not exceed 30°C) for 1 hour. The slices rehydrated during this time. If the gel slices appeared white or opaque after one hour, an additional 10–20 µl of trypsin was added, and they were incubated for another hour at room temperature. Sufficient digestion buffer (40 mM NH_4_HCO_3_/10% ACN) to completely cover the gel slices was added, and the tubes were capped tightly to avoid evaporation. The samples were then incubated overnight at 37°C. Next, the gel slice digests were incubated with 150 µl of pure water for 10 minutes with frequent vortex mixing. The liquid was removed and saved in a new microcentrifuge tube. The gel slice digests were extracted twice with 50 µl of 50% ACN/5% TFA (with mixing) for 60 minutes each time at room temperature. All extracts were pooled and dried in a Speed Vac^®^ at room temperature for 2–4 hours. The extracted peptides were purified and concentrated using ZipTip^®^ pipette tips (Millipore Corporation, Cat# ZTC18S096, Darmstadt, Germany). The peptides eluted from the ZipTip^®^ tips were then ready for mass spectrometric analysis.

### Filter-aided protein digestion

Proteins were digested with trypsin (Trypsin Gold, Mass Spec Grade, Promega, Fitchburg, Wisconsin, USA) using filter-aided sample preparation methods [22]. After proteins were loaded into a 10-kDa filter unit (Pall, Port Washington, New York, USA), UA buffer (8 M urea in 0.1 M Tris-HCl, pH 8.5) and NH4HCO3 (25 mM) were added successively, and the tube was centrifuged at 12,000×g for 30 min at 18°C. Proteins were denatured by incubation with 20 mM DTT at 37°C for 1 h and then alkylated with 50 mM iodoacetamide (IAA) for 45 min in the dark. After the samples had been centrifuged with UA twice and NH4HCO3 four times, the proteins were redissolved in an NH4HCO3 solution and digested with trypsin (1:50) at 37°C overnight. The tryptic peptides were desalted using Oasis HLB cartridges (Waters, Milford, Massachusetts, USA), and the desalted peptides were dried by vacuum evaporation (Thermo Fisher Scientific, Bremen, Germany).

### Peptide tandem mass tag (TMT) labeling

The filter-aided digested peptides were solubilized in 100 mM tetraethylammonium bromide (TEAB) and labeled with 6-plex Tandem Mass Tag Label Reagents provided by Thermo Fisher Scientific (Pierce, Rockford, IL, USA), which had been equilibrated to room temperature immediately before use. Then, 41 μl of anhydrous acetonitrile was added to each tube, and the reagent was allowed to dissolve for 5 min with occasional vortexing. The samples were briefly centrifuged to gather the solution, and 20 µl of the TMT Label Reagent was added. The reaction was then incubated for 2 h at room temperature. After the peptides were labeled with isobaric tags, they were mixed at a 1:1:1:1:1:1 ratio based on the amount of total peptide, which was determined by running equal volumes of labeled samples through liquid chromatography coupled with tandem mass spectrometry (LC-MS/MS) and comparing the total signal intensities of all peptides. Finally, samples from the wild-type group (n=3) and AD group (n=3) were analyzed by two-dimensional LC-MS/MS (2DLC-MS/MS).

### HPLC separation

The TMT-labeled samples were fractionated using a high-pH reversed-phase liquid chromatography (RPLC) column from Waters (4.6 mm × 250 mm, Xbridge C18, 3 µm) and loaded onto the column in buffer A1 (H_2_O, pH=10). The elution gradient progressed from 5–25% buffer B1 (90% ACN, pH=10; flow rate=1 ml/min) over 60 min. The eluted peptides were collected at one fraction per minute. The 60 fractions were dried, resuspended in 0.1% formic acid and pooled into 30 samples by combining fractions 1 and 31, 2 and 32, and so on. The odd-numbered fractions were chosen for further analysis. A total of 15 fractions from urinary peptide mixtures were analyzed by LC-MS/MS.

### LC-MS/MS analysis

The in-gel digested peptides and TMT-labeled peptides were analyzed with a reverse-phase C18 self-packed capillary LC column (75 µm × 100 mm). The elution gradient progressed from 5–30% buffer B2 (0.1% formic acid, 99.9% ACN; flow rate=0.3 µL/min) over 40 min. A Triple TOF 5600 mass spectrometer was used to analyze the peptides eluted from LC, and each fraction was run twice. MS data were acquired using the high-sensitivity mode with the following parameters: 30 data-dependent MS/MS scans per full scan, full scans acquired at a resolution of 40,000, MS/MS scans acquired at a resolution of 20,000, rolling collision energy, charge state screening (including precursors with a charge state of +2 to +4), dynamic exclusion (exclusion duration 15 s), an MS/MS scan range of 100-1800 m/z, and a scan time of 100 ms.

### Data analysis

The MS/MS data were analyzed with Mascot software (version 2.4.1, Matrix Science, London, UK), and proteins were identified by comparing the peptide spectra against the Swissprot_2014_07 databases (taxonomy: mus). Trypsin was selected as the digestion enzyme, up to two missed cleavage sites were allowed, and carbamidomethylation of a cysteine was defined as a fixed modification. The precursor ion mass tolerance and the fragment ion mass tolerance were 0.05 Da. The TMT-labeled protein identification Mascot results were validated by using Scaffold Proteome Software (version Scaffold_4.3.3, Proteome Software Inc., Portland, OR). Peptide identification was accepted at a false discovery rate (FDR) of less than 1.0% at the protein level and if the sample produced at least 2 unique peptides. The Scaffold Q+ was used for Label-Based Quantification (TMT, iTRAQ, SILAC, etc.) of peptides and proteins. Normalized reporter ion intensities were used to calculate the relative protein abundance. The protein ratios were then quantified by the median of the transformed reporter ion intensity ratios [23]. Label-free quantification was performed using Progenesis LC-MS software (version 4.1, Nonlinear, UK) as described previously [24].

### Statistical analysis

All statistical analysis was performed with the Statistical Package for Social Studies software (SPSS, version 16, IBM). Comparisons between independent groups were conducted using one-way ANOVA followed by post hoc analysis with the least significant difference (LSD) test, and P-values less than 0.05 were considered to indicate statistically significant differences.

## Results and discussion

### Urinary proteome changes in APP (swe)/PSEN1^dE9^ transgenic mice

The urinary proteins of 4-month-old mice were digested by an in-gel method and quantified by a label-free method. A total number of 443 protein groups were identified in the urinary proteome at an FDR < 1%, and each generated at least 2 unique peptides. The levels of 33 urinary proteins were significantly different between the AD and wild-type groups (Table 1); among these proteins, cadherin-16 levels were different in both the >85 kDa and 50-85 kDa lanes, sulfhydryl oxidase 1 levels were different in both the 50-80 kDa and 30-50 kDa lanes, and deoxyribonuclease-1 levels were different in both the 15-30 kDa and <15 kDa lanes. The fold change of the significantly changed protein levels was more than 1.5. In these 33 differential proteins, 17 had been reported to be associated with AD. Specifically, 15 had been reported to be associated with the pathology mechanisms of AD, while 9 were identified as direct AD biomarkers (Table 1).

**Table 1.**
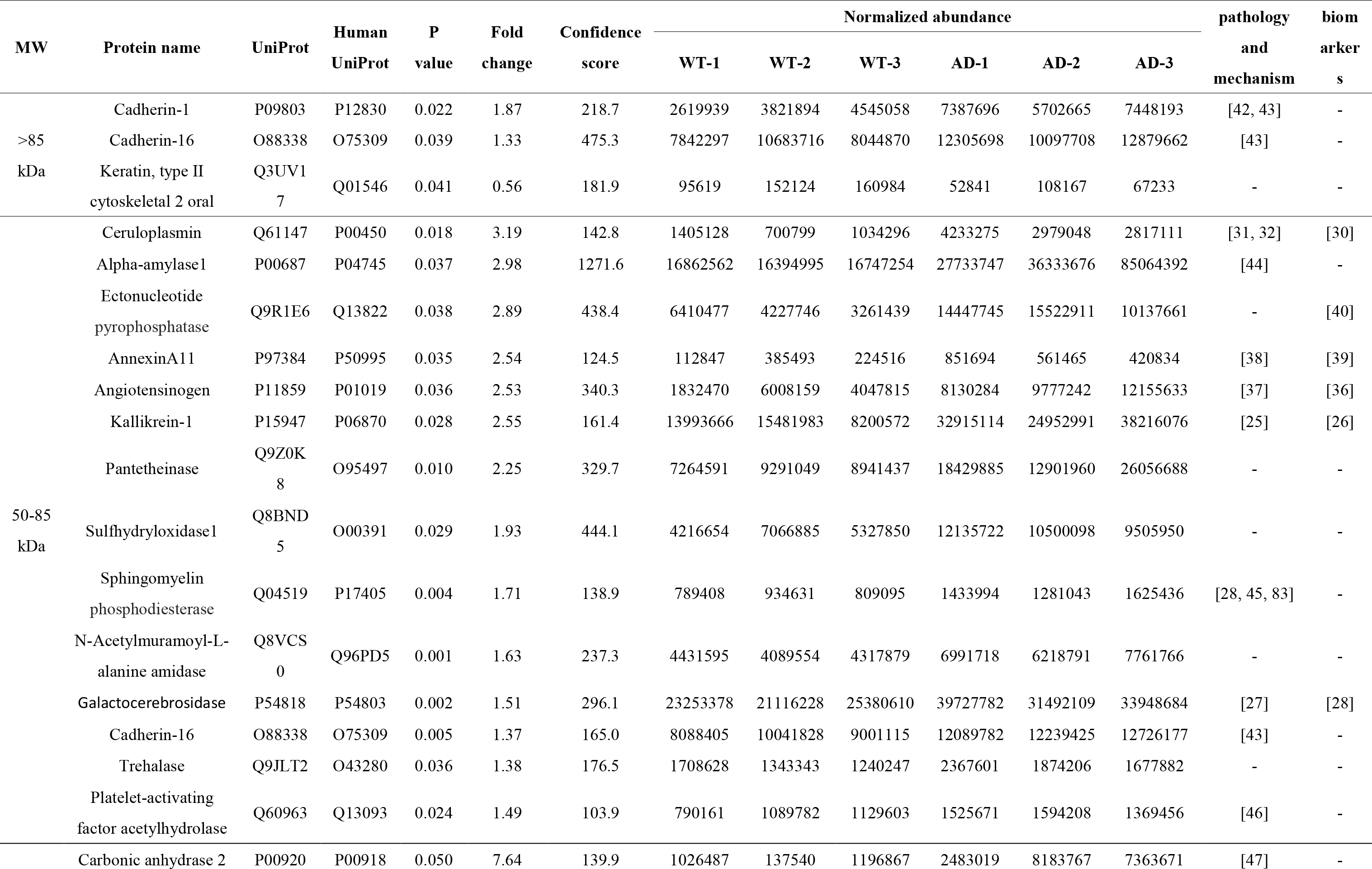
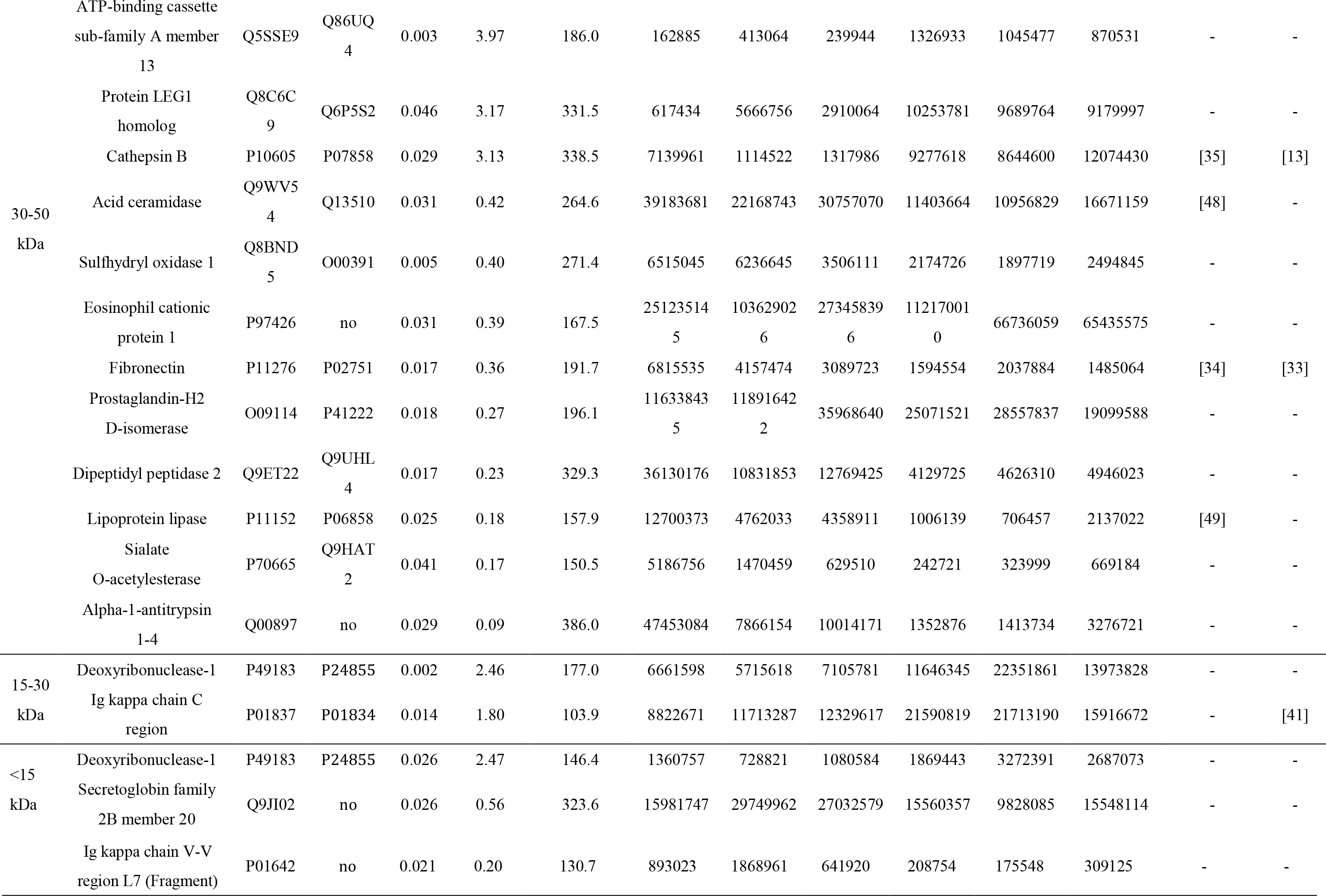
Details of differential urinary proteins of 4-month-old mice

Seven proteins relate to the development of AD as well as biomarkers of AD. (1) Kallikrein-1 (KLK1) levels were different in 4-, 6- and 8-month-old groups than the wild-type group, and the kallikrein-kinin system mediates inflammation in AD in vivo[25]. Kallikrein-6, one member of the kallikrein family, was reported as a biomarker of AD and to have increased levels in CSF and serum and decreased levels in tissue [26]. (2) Galactocerebrosidase (GALC) levels were different in 4- and 8-month-old groups than the wild-type group, and a deficiency in galactosylceramidase potentially contributed to neurodegenerative disease [27]. Ceramide has been suggested to participate in the neuronal cell death that leads to AD. GALC, as a gene connected to ceramide metabolism, was upregulated in the brain tissue of AD patients, making it an attractive candidate for diagnostic purposes and for intervening in neurodegenerative processes [28]. (3) Ceruloplasmin (CERU) levels were higher in both 4- and 8-month-old groups than in the control group, with the same trend of increased levels found in the serum of AD patients[29]. The levels of CSF diagnostic markers, such as Aβ42, tau and phospho-tau, were correlated with lower 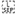 plasma copper and CERU levels in patients with Alzheimer’s disease[30]. The ratio of CERU concentrations measured by enzymatic methods (eCP) to those measured by immunological methods (iCP), eCP/iCP, reflects the high specificity of AD patients as well as a decreased risk of having AD[31]. CERU had less ferroxidase activity in AD patients than wild-type patients, which contributed to the development of AD[32]. (4) Fibronectin (FINC) levels were different in 4- and 8-month-old groups than in the wild-type group and were reported to be a novel biomarker for AD from blood[33]. In addition, FINC levels were significantly lower in plasma in mild cognitive impairment 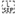 (MCI) patients than healthy patients, which provides further insight into the biological pathways and processes that underpin the pathophysiology and progression of MCI and AD[34]. (5) Cathepsin B (CATB) levels were different in 4- and 8-month-old groups than the wild-type group. CATB has been previously reported to be upregulated in brain tissues from APP/PS1 transgenic mice, and its levels changed in the same direction relative to healthy mice as they changed in the serum of AD patients relative to control patients, making CATB a potential biomarker of AD[13]. In addition, CATB produces brain pyroglutamate amyloid beta, which represents a potential AD therapeutic[35]. (6) Angiotensinogen (ANGT) was upregulated in 4-month-old groups and has exhibited increased levels in the CSF of AD patients[36]. As a component of the renin-ANGT system (RAS), ANGT is helpful for AD processes[37]. (7) Annexin A11 (ANX11) was upregulated in 4-month-old groups. Annexin A1 was reported to be expressed strongly in the microglia of AD patients, which aids the surveillance of microglia and the maintenance of brain homeostasis by using Annexin A1-dependent mechanisms[38], and Annexin A5 was reported to be a biomarker of AD and present at increased levels in the plasma of AD patients[39]. The protein-normalized abundances and spectral counts of all seven differential proteins were greater in every mouse in the high-abundance group than those in the low-abundance group.

Two other differential proteins have been mentioned as potential AD biomarkers. (1) Ectonucleotide pyrophosphatase/phosphodiesterase family member 2 (ENPP2) levels were higher in the CSF of AD patients than the CSF of healthy controls. ENPP2 can be used to specifically discriminate AD from Lewy body dementia, making it a candidate AD biomarker[40]. (2) The Ig kappa chain C (IGKC) region was reported to be a potential serum biomarker of AD at the early stage[41].

Eight differential proteins are associated with the mechanism of AD pathology. (1) CDH1, the gene of cadherin-1, is an AD risk gene that has significant association in Caribbean Hispanic individuals[42]. (2) As glycosylated transmembrane proteins, cadherins such as cadherin-1 and cadherin-16 also directly interact with the presenilin-1/g-secretase complex, which contributes to the modulation of Aβ peptide formation and thus plays a critical role in AD etiology[43]. (3) Alpha-amylase 1 (AMY1) reflects M3 muscarinic agonist activity in AD[44]. (4) ASMase, the gene of sphingomyelin phosphodiesterase, which is connected to ceramide metabolism, was upregulated in the brain tissue of AD patients, making it an attractive candidate for both diagnostic purposes and intervening in neurodegenerative processes[28]. Significant correlations were observed between brain sphingomyelin phosphodiesterase levels and the levels of amyloid beta peptide (Aβ) and phosphorylated tau protein, which contribute to AD disease pathogenesis[45]. (5) Platelet-activating factor acetylhydrolase (PAFA) was present in higher levels in the plasma of AD patients than that of healthy controls, and these higher levels were associated with oxidative damage of low-density lipoproteins (ox-LDL), which contributes to the inflammation and oxidative stress of plasma lipoproteins and is strongly associated with Alzheimer’s disease[46]. (6) Plasma carbonic anhydrase 2 (CAH2) was elevated in AD patients and may thus play a role in the pathogenesis of AD[47]. (7) Acid ceramidase (ASAH1) levels were elevated in AD brain, suggesting that acid ceramidase might play a role in controlling neuronal apoptosis and that acid ceramidase-mediated signaling pathways might be involved in the molecular mechanism of AD[48]. (8) Lipoprotein lipase (LIPL) is associated with neurite pathology, and its levels were reduced in the dentate gyrus of AD brains[49]. Some differential proteins have not been reported in research papers and may provide clues to candidates involved in the AD pathology mechanism.

The urinary proteins of 6-month-old mice were digested by a filter-aided method and quantified by a TMT-labeling method. At the protein level, 310 protein groups were identified in the urinary proteome at an FDR < 1%, and each generated at least 2 unique peptides. The levels of 33 urinary proteins were significantly different between the AD and wild-type groups, and their fold changes were greater than 1.5. Eight of these 33 differential proteins had been reported to be associated with AD. To be specific, 6 were associated with the pathology mechanisms of AD, while 3 had been directly referred to be AD biomarkers (Table 2).

**Table 2.** Details of differential urinary proteins of 6-month-old mice

| Protein name | UniProt | Human | P | Fold | Spectral | MW | Normalized ratio |  |  |  |  |  | pathology and<br>mechanism | biomarker |
| --- | --- | --- | --- | --- | --- | --- | --- | --- | --- | --- | --- | --- | --- | --- |
|  |  | UniProt | value | change | count |  | WT-1 | WT-2 | WT-3 | AD-1 | AD-2 | AD-3 |  |  |
| Major urinary protein 1 | P11588 | NO | 0.004 | 10.92 | 1657 | 21 kDa | 0.45 | 0.95 | 1.6 | 8.45 | 13.9 | 10.4 | - | - |
| Neogenin | P97798 | Q92859 | 0.005 | 7.22 | 3 | 38 kDa | 1 | 1.45 | 0.1 | 4.8 | 7.75 | 5.85 | [51] | - |
| CMRF35-like molecule<br>2 | Q8K249 | Q496F6 | 0.004 | 5.31 | 3 | 22 kDa | 0.5 | 3 | 3.4 | 12.45 | 9.7 | 14.5 | - | - |
| Major urinary protein 5 | P11591 | no | 0.011 | 3.79 | 326 | 21 kDa | 1.05 | 4 | 3 | 12.45 | 7.55 | 10.5 | - | - |
| WAP four-disulfide<br>core domain protein 12 | Q9JHY<br>3 | Q8WWY7 | 0.029 | 2.94 | 5 | 10 kDa | 0.9 | 0.95 | 0.65 | 3.25 | 2.5 | 1.6 | - | - |
| Keratin, type I<br>cytoskeletal 19 | P19001 | P08727 | 0.014 | 2.41 | 9 | 45 kDa | 1 | 1.2 | 1.25 | 3.35 | 2.05 | 2.9 | - | - |
| Cadherin-15 | P33146 | P55291 | 0.015 | 2.06 | 3 | 86 kDa | 1 | 0.6 | 0.75 | 1.95 | 1.4 | 1.5 | - | - |
| Charged multivesicular<br>body protein 2a | Q9DB3<br>4 | O43633 | 0.027 | 1.88 | 4 | 25 kDa | 1 | 0.8 | 0.65 | 1.35 | 1.35 | 1.9 | [52] | - |
| Cadherin-11 | P55288 | P55287 | 0.024 | 1.87 | 2 | 88 kDa | 1 | 0.7 | 0.55 | 1.65 | 1.25 | 1.3 | - | - |
| Peroxisomal carnitine | Q9DC5<br>0 | Q9UKG9 | 0.030 | 1.86 | 2 | 70 kDa | 1 | 1.05 | 0.85 | 2.25 | 1.75 | 1.4 | [53] | - |
| Desmoplakin | E9Q557 | P15924 | 0.041 | 1.80 | 27 | 333 kDa | 0.8 | 0.85 | 0.8 | 1.3 | 1.2 | 1.9 | - | - |
| Pancreatic secretory<br>granule membrane<br>major glycoprotein GP2 | Q9D733 | P55259 | 0.048 | 1.79 | 3 | 59 kDa | 1 | 0.8 | 0.55 | 1.55 | 1.05 | 1.6 | - | - |
| Alpha-1-antitrypsin 1-4 | Q00897 | no | 0.022 | 1.75 | 6 | 46 kDa | 0.9 | 0.8 | 0.9 | 1.45 | 1.85 | 1.25 | - | - |
| Heat shock cognate 71 kDa protein | P63017 | P11142 | 0.020 | 1.67 | 7 | 71 kDa | 0.9 | 0.9 | 0.95 | 1.45 | 1.3 | 1.85 | [54] | - |
| Kynurenine/alpha-aminoadipate aminotransferase, mitochondrial | Q9WV M8 | Q8N5Z0 | 0.002 | 1.63 | 2 | 48 kDa | 1 | 1.15 | 0.85 | 1.65 | 1.6 | 1.65 | - | - |
| Semaphorin-4A | Q62178 | Q9H3S1 | 0.002 | 1.62 | 5 | 83 kDa | 0.85 | 0.7 | 0.7 | 1.2 | 1.15 | 1.3 | - | - |
| Serine protease inhibitor A3M | Q03734 | no | 0.006 | 1.53 | 3 | 47 kDa | 0.9 | 0.95 | 0.8 | 1.2 | 1.45 | 1.4 | - | - |
| Leukemia inhibitory factor receptor | P42703 | P42702 | 0.044 | 1.51 | 6 | 122 kDa | 0.9 | 0.9 | 1.05 | 1.45 | 1.15 | 1.7 | [55] | - |
| Kallikrein-1 | P15947 | P06870 | 0.028 | 0.61 | 79 | 29 kDa | 1 | 1.5 | 1.35 | 0.8 | 0.75 | 0.8 | [25] | [26] |
| Prominin-1 | O54990 | O43490 | 0.027 | 0.60 | 2 | 97 kDa | 1 | 0.75 | 0.75 | 0.55 | 0.4 | 0.55 | - | - |
| Glycerophosphodiester phosphodiesterase domain-containing protein 3 | Q99LY 2 | Q7L5L3 | 0.039 | 0.59 | 7 | 38 kDa | 0.9 | 1.35 | 1.4 | 0.65 | 0.7 | 0.8 | - | - |
| Ig kappa chain C region | P01837 | P01834 | 0.003 | 0.57 | 25 | 12 kDa | 1 | 0.85 | 0.8 | 0.5 | 0.5 | 0.5 | - | [41] |
| Solute carrier family 12 member 3 | P59158 | P55017 | 0.030 | 0.55 | 3 | 111 kDa | 1 | 0.9 | 1.3 | 0.5 | 0.75 | 0.5 | - | - |
| Meprin A subunit beta | Q61847 | Q16820 | 0.029 | 0.54 | 6 | 79 kDa | 1.1 | 1.05 | 0.9 | 0.3 | 0.7 | 0.65 | - | - |
| Solute carrier family 23 member 1 | Q9Z2J0 | Q9UHI7 | 0.014 | 0.54 | 9 | 66 kDa | 1 | 0.7 | 0.8 | 0.4 | 0.45 | 0.5 | - | - |
| Sushi domain-containing protein 2 | Q9DBX 3 | no | 0.001 | 0.54 | 3 | 90 kDa | 0.9 | 1 | 0.9 | 0.5 | 0.45 | 0.55 | - | - |
| Villin-1 | Q62468 | P09327 | 0.011 | 0.52 | 8 | 93 kDa | 1 | 0.7 | 0.8 | 0.4 | 0.45 | 0.45 | - | - |
| Ezrin | P26040 | P15311 | 0.027 | 0.51 | 11 | 69 kDa | 0.9 | 0.55 | 0.7 | 0.35 | 0.4 | 0.35 | - | - |
| Chloride intracellular channel protein 4 | Q9QYB1 | Q9Y696 | 0.008 | 0.50 | 3 | 29 kDa | 1 | 0.7 | 0.9 | 0.45 | 0.45 | 0.4 | - | - |
| Na(+)/H(+) exchange regulatory cofactor NHE-RF3 | Q9JIL4 | Q5T2W1 | 0.013 | 0.48 | 4 | 56 kDa | 1 | 0.8 | 0.7 | 0.3 | 0.45 | 0.45 | - | - |
| Ras-related protein Rab-10 | P61027 | P61026 | 0.041 | 0.43 | 2 | 23 kDa | 1 | 1.05 | 1.65 | 0.4 | 0.75 | 0.45 | - | [50] |
| Na(+)/H(+) exchange regulatory cofactor NHE-RF1 | P70441 | O14745 | 0.011 | 0.42 | 13 | 22 kDa | 1 | 0.65 | 0.75 | 0.3 | 0.35 | 0.35 | - | - |
| Ig kappa chain V-V region L7 (Fragment) | P01642 | no | 0.005 | 0.38 | 10 | 13 kDa | 1 | 0.8 | 0.7 | 0.35 | 0.3 | 0.3 | - | - |

Three differential proteins (KLK1, IGKC, and RAB10) have been suggested as AD biomarkers. The target gene RAB10, which is regulated by miRNA-369-3p, miRNA-30e-5p, miRNA-30e-3p, and miRNA-655, may play key roles in the progression and development of AD and may be a biomarker of AD[50].

Six differential proteins are associated with the mechanism of AD pathology. (1) Neogenin (NEO1) expresses weak NEO1 immunoreactivity in a small subset of amyloid plaques of AD brains[51]. (2) Charged multivesicular body protein 2a (CHM2A) levels were elevated in the 6-month-old group. Mutations in charged multivesicular body protein 2B (CHMP2B) can give rise to FTLD-UPS[52]. (3) Peroxisomal carnitine (OCTC) levels were higher in the 6-month-old group than the wild-type group, and new therapies acting on peroxisomal metabolism might be developed to prevent cognitive decline and other age-related neurological disorders[53]. (4) Heat shock cognate 71 kDa protein (HSP7C) was upregulated in 6-month-old groups, while its gene level was significantly lower across the three brain regions in AD patients than in controls patients, suggesting 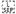 their participation in AD pathogenesis[54]. (5) Cooperative expression of leukemia inhibitory factor (LIF) and its receptor (LIFR) in AD hippocampus may indicate a role for LIF in neuronal damage or repair at these sites[55].

The urinary proteins of 8-month-old mice were digested by an in-gel method and quantified by a label-free method. At the protein level, 273 proteins were identified in the urinary proteome at an FDR < 1%, and each generated at least 2 unique peptides. The levels of 89 urinary proteins were significantly different between the AD and wild-type groups. Thirty-three of these 89 differential proteins have been associated with AD. Specifically, 29 are associated with the mechanism of AD pathology, while 17 have been suggested as candidate AD biomarkers. (Table 3)

**Table 3.**
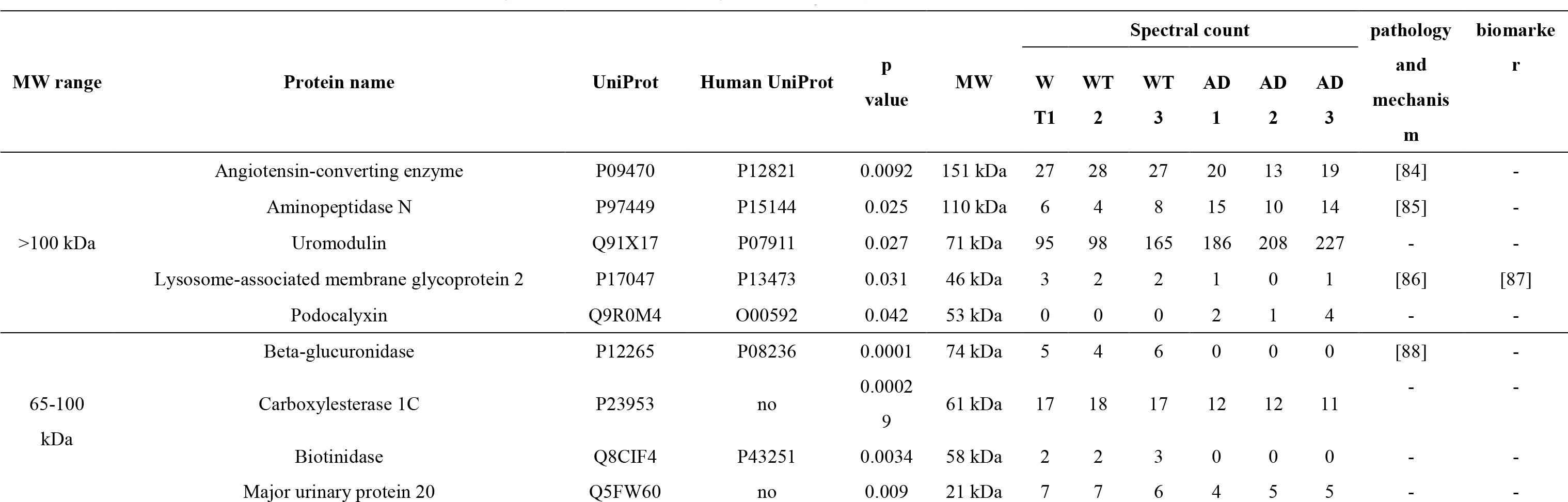
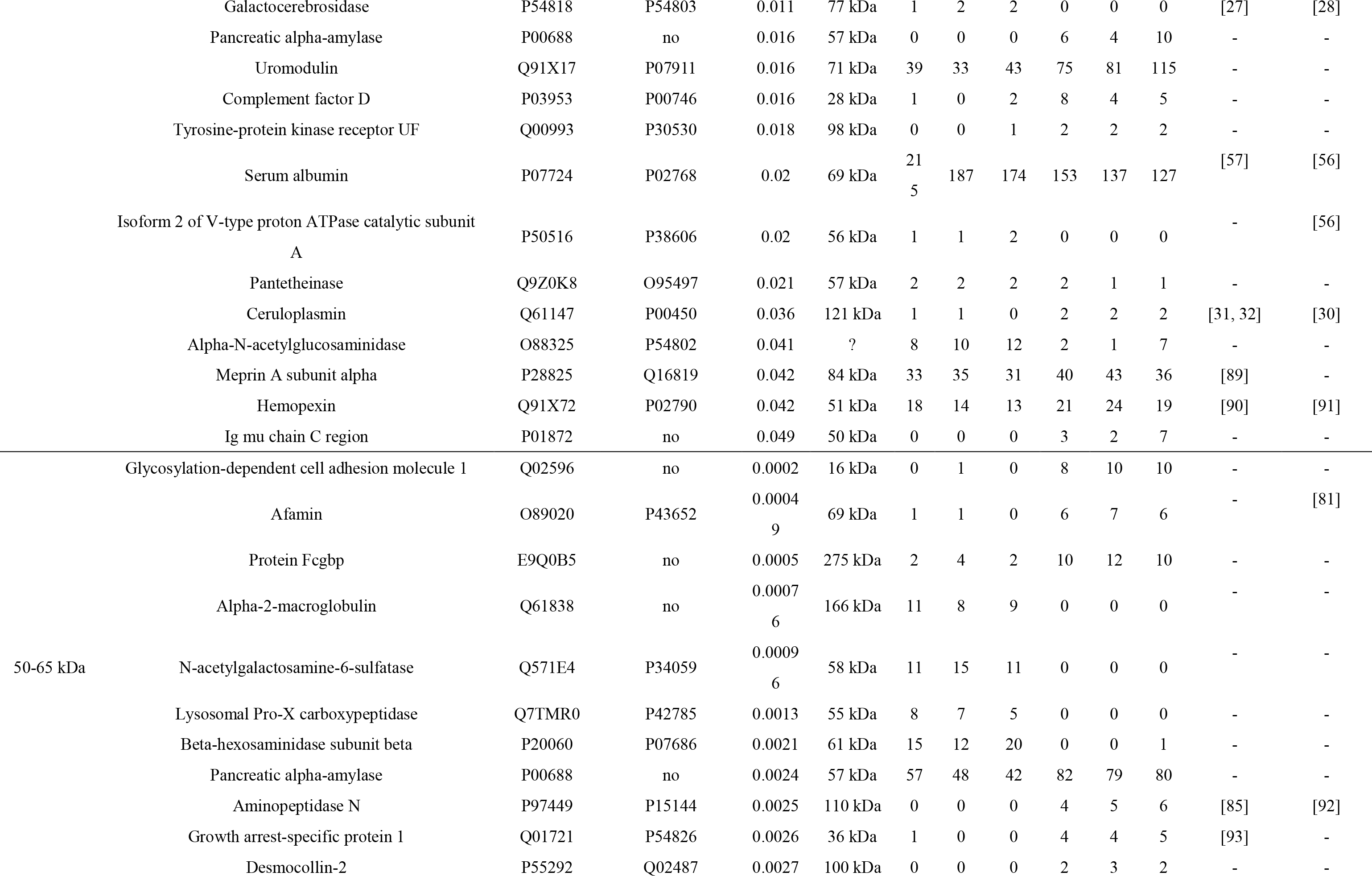
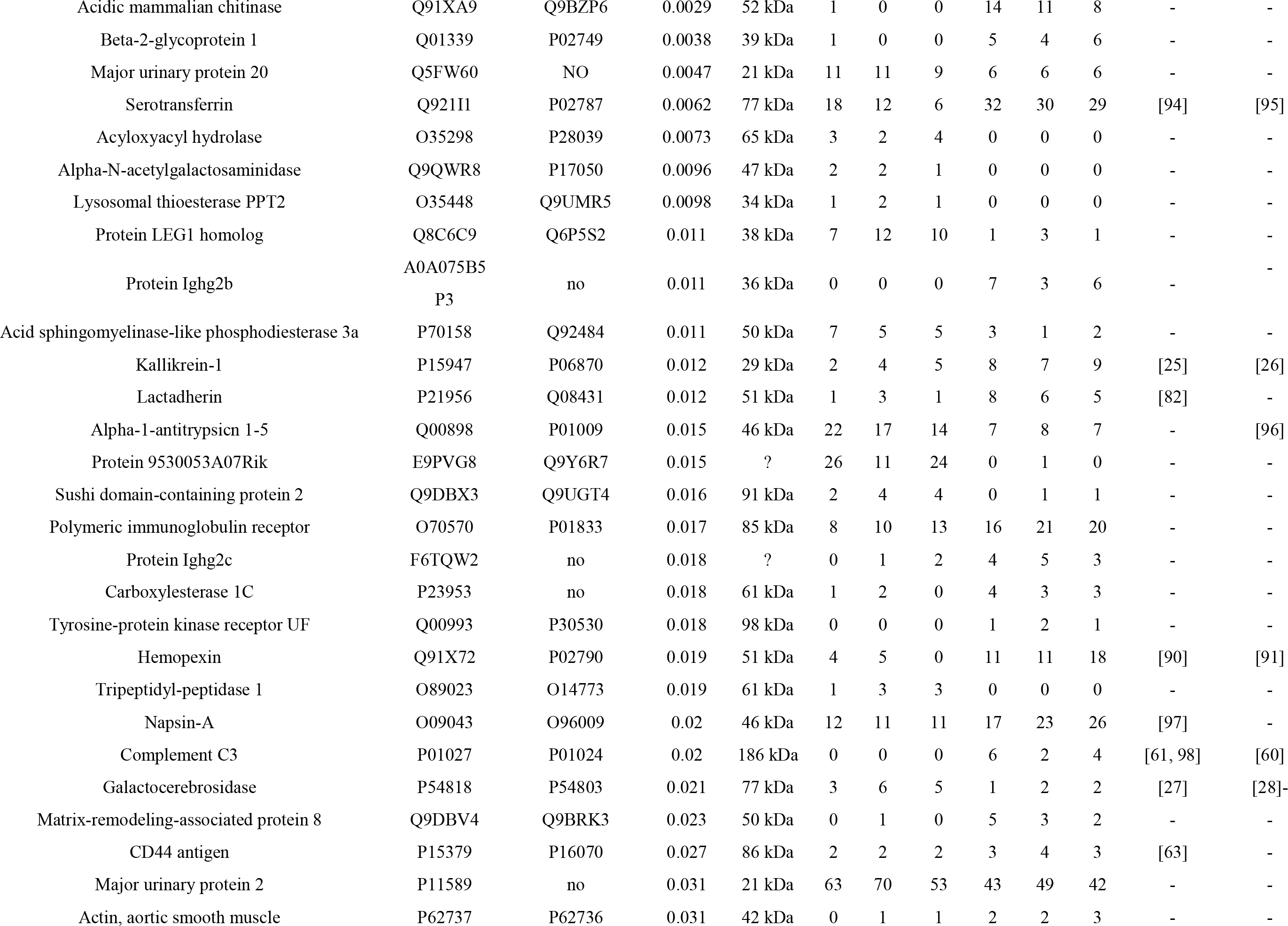
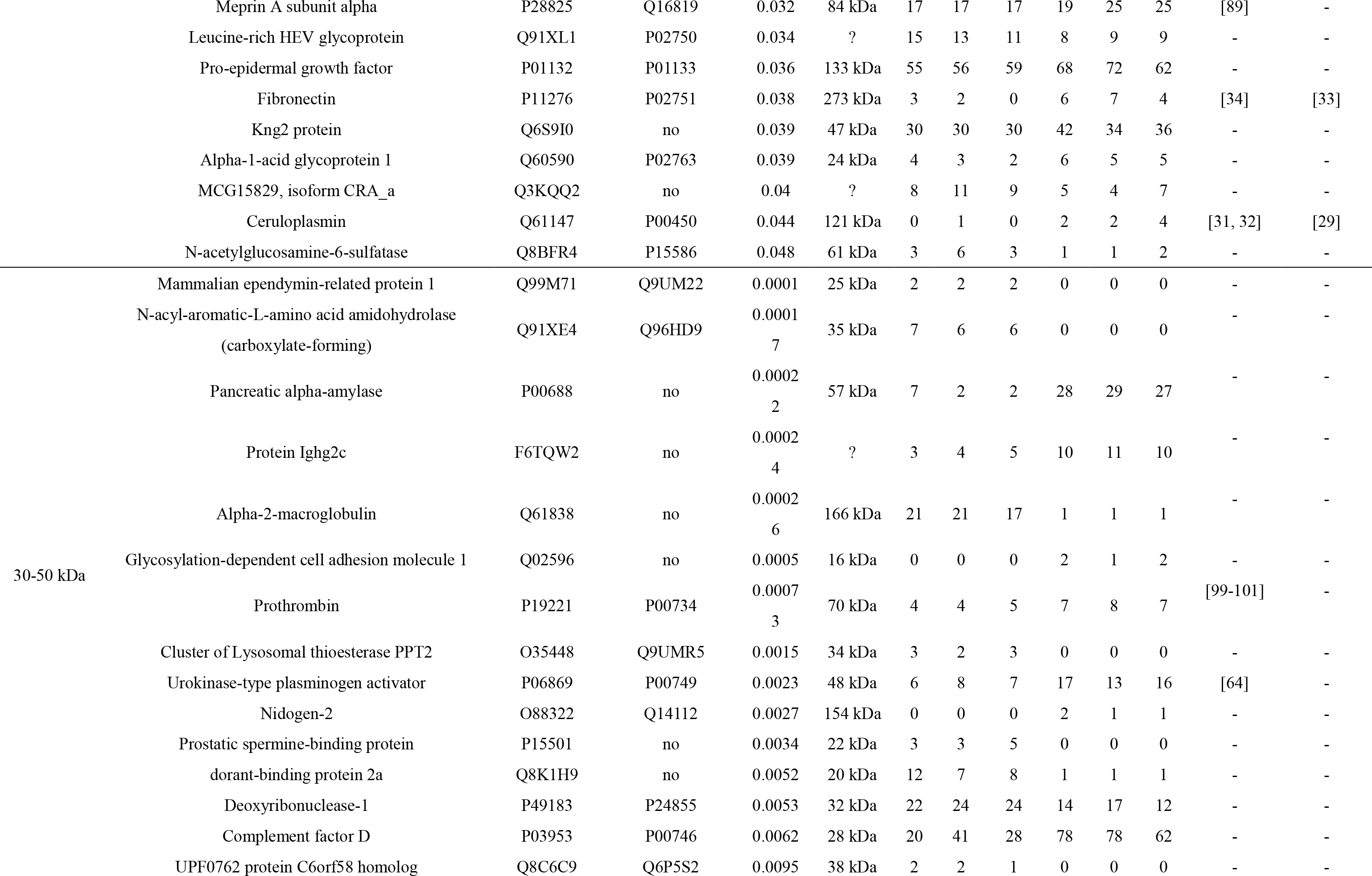
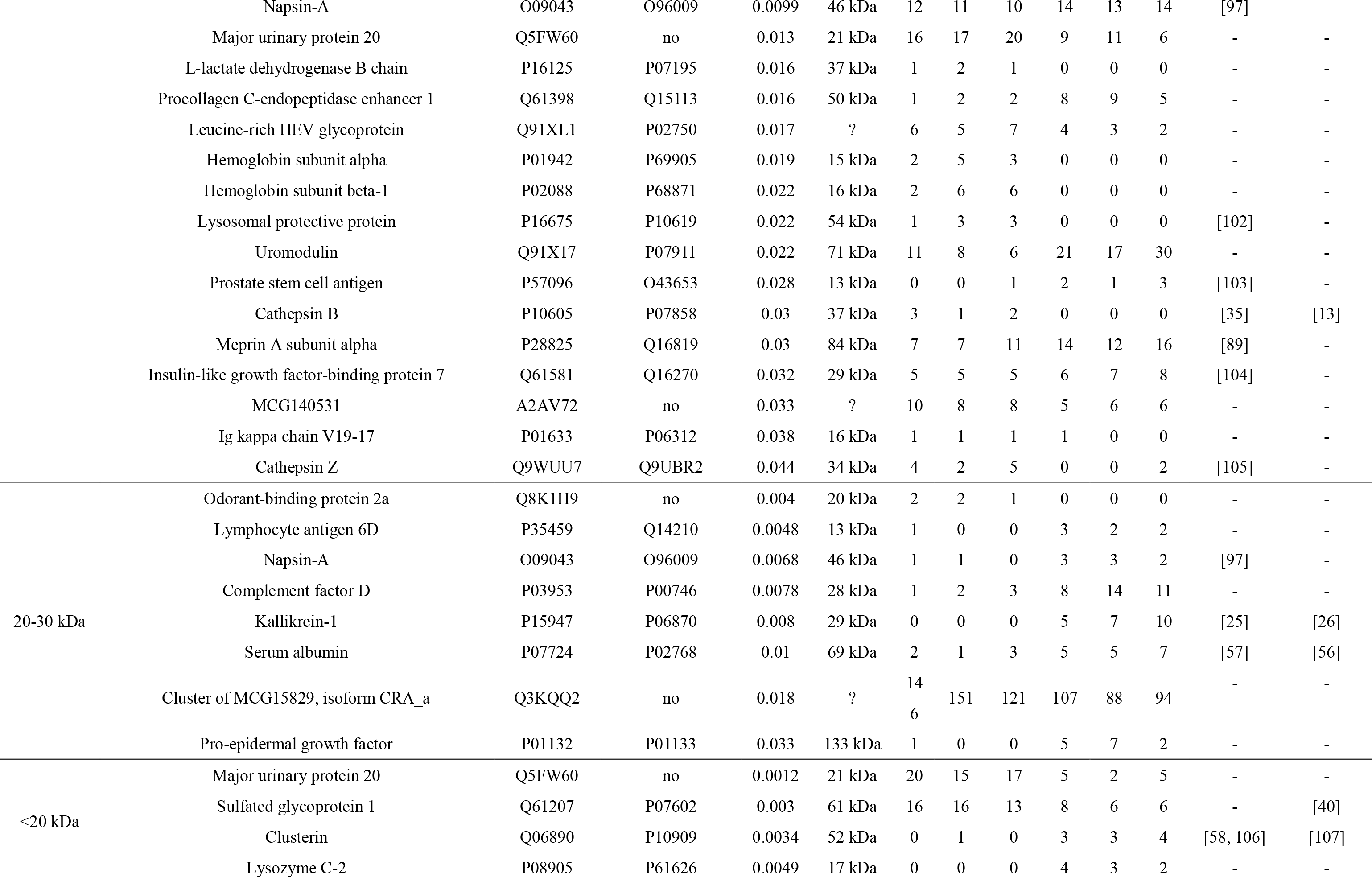
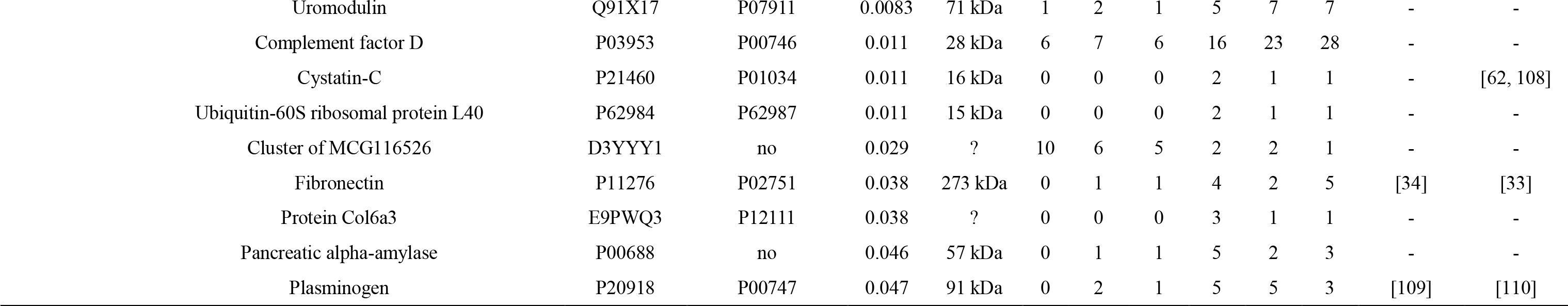
Details of differential urinary proteins of 8-month-old mice

Thirteen proteins have been previously reported to be related to the development of AD as well as biomarkers of AD, while 5 of them (KLK1, GALC, CERU, FINC, and CATB) had been already noted in 4-month-old groups. For example, (1) Serum albumin (ALBU) was differentially expressed between brains of AD mouse models and those of healthy mice[56], and serum levels of albumin-amyloid beta complexes were lower in AD patients than healthy controls, which is very useful to apply in monitoring the progression of AD[57]. (2) Clusterin (CLUS) levels were higher in AD patients than controls, especially in regions with most abundant in Aβ, facilitating the development of AD[58]. As a heterodimeric glycoprotein, clusterin was more abundant in the CSF of some neurodegenerative disease patients than that of healthy controls and could thus serve as a potential biomarker to differentiate Parkinson’s disease (PD) from dementia with Lewy bodies (DLB)[59]. (3) CSF complement C3 (CO3) is a staging biomarker in AD[60]. In addition, the dysregulation of neuron-glia interactions through NFκB/ C3/C3aR signaling may contribute to synaptic dysfunction in AD[61]. Seventeen differential proteins have been mentioned as potential AD biomarkers. For example, (1) sulfated glycoprotein 1 (SAP) is a novel CSF biomarker for neurodegeneration and has been validated by a high-throughput multiplexed targeted proteomic assay[40]. (2) Cystatin-C (CYTC) is a novel CSF biomarker for staging early AD[62]. Twenty-nine differential proteins are associated with the mechanism of AD pathology. For example, (1) CD44 antigen (CD44) gene expression was greater in lymphocytes derived from AD patients than in those from healthy controls, suggesting that it plays roles in the peripheral immune response during the development of AD[63]. (2) The rs2227564 polymorphism of the urokinase-type plasminogen activator (UROK) gene increased the risk of AD[64]. Other differential proteins are annotated in Table 3.

Functional annotation of differential urinary proteins was performed using DAVID[65]. The differential proteins at three time points were classified to be involved with certain biological processes (Figure 4), molecular components, and molecular functions (Supplemental Figures 1 and 2).

**Figure 4.**
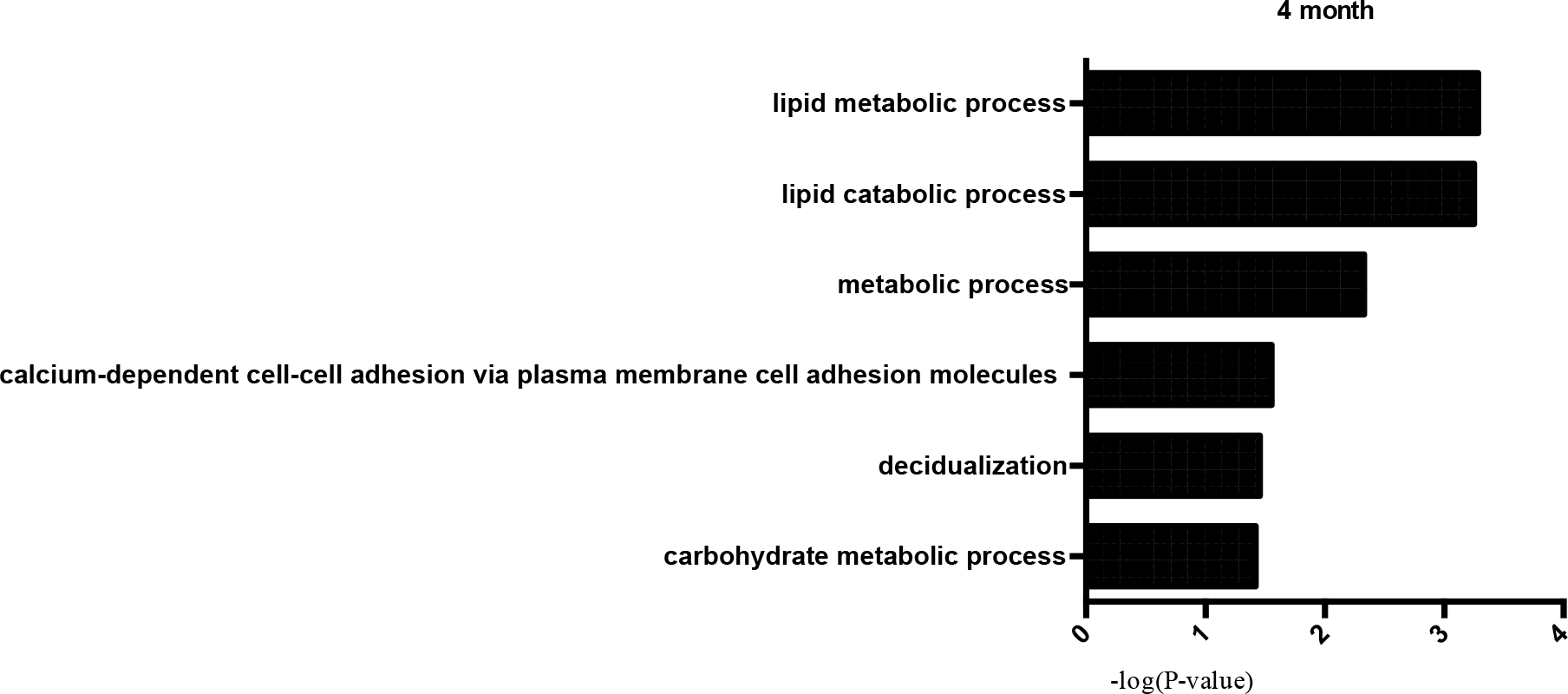
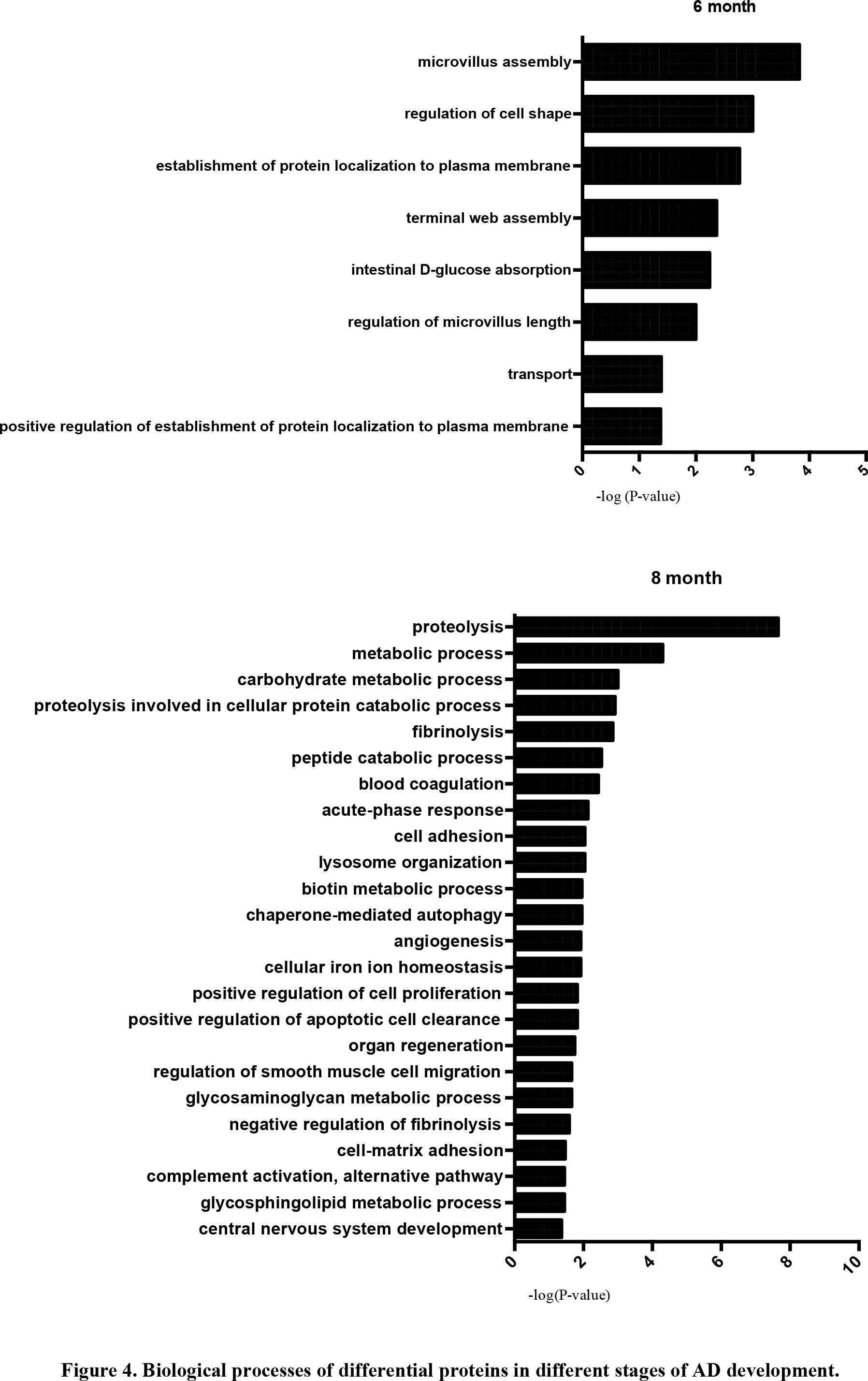
Biological processes of differential proteins in different stages of AD development.

In 4-month-old mice, differential urine proteins involved in lipid metabolism and lipid catabolism were enriched. As lipid metabolism is a fundamental process for brain development and function, aberrant lipid metabolism was not surprisingly common in an animal model of AD that relates to AD pathogenesis[66-68]. For example, AD-induced perturbation of niche fatty acid metabolism can suppress the homeostatic and regenerative functions of neural stem cells, supporting the mechanism of AD pathogenesis[66]. Abnormal lipid metabolism influences Aβ metabolism and deposition in both brain parenchyma and vasculature, as well as tau hyperphosphorylation and aggregation, which then likely triggers a series of downstream catalytic events that eventually affect the progression of AD pathogenesis[67]. More importantly, lipid rafts from human cerebral cortex are associated with the pathogenesis of early AD, as β-secretase/AβPP (amyloid-β protein precursor) interactions and lipid raft microviscosity are strongly and positively correlated in AD frontal and entorhinal cortices, indicating that the aberrant lipid metabolism had already occurred in the early stage of AD[69]. Decidualization and calcium-dependent cell-cell adhesion via plasma membrane cell adhesion molecules, which are not reported in AD studies, may play roles in the mechanism of AD pathology. Recent studies demonstrated that using a triple receptor agonist (TA), which activates GIP-1, GIP and glucagon receptors at the same time, reduces the total amount of beta amyloid in an APP/PS1 transgenic mouse model through a 2-month TA treatment [70]. However, this research used 6-month-old transgenic mice for a 2-month treatment, and beta amyloid had already appeared. We thus suppose that a more effective treatment can be achieved when the 4-month-old transgenic mice are used for therapy.

The differential proteins in 6-month-old mice were involved in transport processes, while axonal transport (AT) defects play an important role in the pathogenesis of AD[71]. Other biological processes such as microvillus assembly, the regulation of cell shape and protein localization to the plasma membrane may contribute to the formation of beta amyloid plaque deposits in the brain of AD patients, which provides potential clues for the studying the mechanism of AD.

A larger number of differential urine proteins were identified in 8-month-old mice and indicated other AD-related biological processes. Proteolysis processes were overrepresented and reported to be associated with APP proteolysis, which is consistent with the pathology of AD[72]. In addition, the urinary proteins that were related to fibrinolysis may be factors contributing to AD[73]. What’s more, lysosome organization accumulated at amyloid plaques in mouse models of AD, which gave rise to potential therapies of AD patients[74]. Lastly, angiogenesis processes were reported to contribute to the pathogenesis of AD, as they are a common feature of amyloid beta (Aβ) plaques[75, 76], and the levels of some angiogenesis factors were elevated in the plasma of AD patients[77].

In summary, in the biological process category, metabolic and carbohydrate metabolic processes were overrepresented in 4-and 8-month-old AD mice. The differential urine proteins in 4- and 8-month-old mice indicated an enrichment of metabolic process, while the three subtypes of AD can be distinguished by metabolic profiling[78]. Considering that the biological processes whose activities changed in this experiment did not extensively overlap at these three time points, this result indicates that only three different stages might occur, implying that different therapeutic strategies may be appropriate for different stages. If biomarkers can be found in human as early as was found for the 4-month-old in this model, we may have an early intervention window in which disease development can be stopped.

In the cellular component category (Supplemental Figure 1), most differential proteins came from the extracellular exosome, the extracellular space, the anchored component of membranes, and blood microparticles, whereas a small number of differential proteins were derived from organelles such as the lysosome and the Golgi apparatus. In the molecular function category (Supplemental Figure 2), hydrolase activity was overrepresented in 4- and 8-month-old mice; calcium ion binding was overrepresented in 4-month-old mice; protein complex binding, sodium ion transmembrane transporter activity and protein binding were overrepresented in 6-month-old mice; and peptidase, endopeptidase, serine-type peptidase activities were overrepresented in 8-month-old mice.

### IPA of differential proteins in AD development

To identify the differential proteins with mostly affected cellular death and survival, ingenuity pathway analysis (IPA) was applied to cell death and survival networks (Figure 5). Cadherin-1, which was reported to be associated with AD[42], is a core component of these networks in 4-month-old mice. CDH16, CP, ENPP2, AGT, SMPD1, PLA2G7 were upregulated and play roles in AD[28, 29, 36, 40, 43, 46], whereas sialate O-acetylesterase was downregulated in 4-month-old mice. In 6-month-old mice, NEO1, CROT, GP2, SEMA4A and LIFR were present in higher levels in transgenic mice than the control group, and NEO1 as well as CROT were associated with the pathology and mechanism of AD[51, 79]. In contrast, MEP1B, CLIC4, and PDZK1 levels were lower in transgenic mice than the control group. FINC was the core component of the network in 8-month-old mice and plays roles in AD[80]. In addition, AFM, MFGE8, C3, CD44 and PLAU were upregulated in transgenic mice and are potential biomarkers or play roles in the development of AD[61, 63, 64, 81, 82], while VNN1, HEXB and CTSZ were downregulated in transgenic mice when compared with control groups.

**Figure 5.**
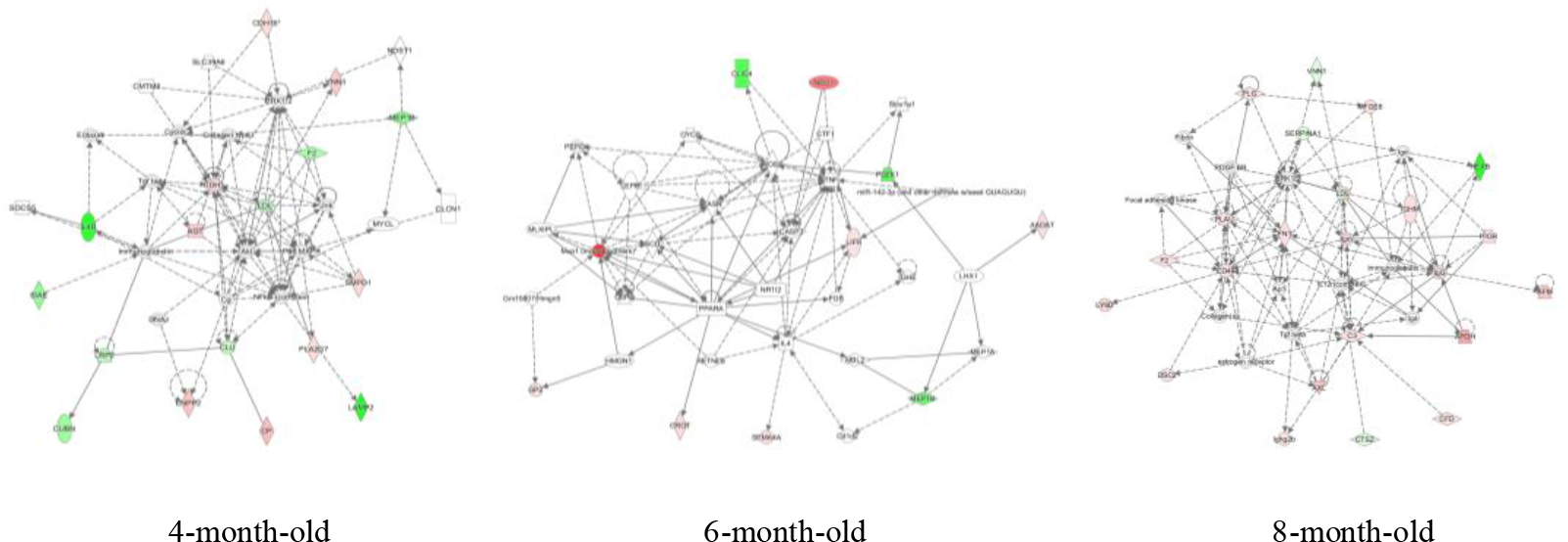
IPA of differential proteins in AD development

## Conclusion

Our results indicated that urine proteins enable AD early detection before beta amyloid plaque deposition, which may provide an opportunity for intervention.

## Acknowledgements

This research was supported by the National Key Research and Development Program of China (2016 YFC 1306300), Key Basic Research Program of the Ministry of Science and Technology of China (2013FY114100), Beijing Natural Science Foundation (7173264, 7172076), Beijing cooperative construction project (110651103) Beijing Normal University (11100704) Peking Union Medical College Hospital (2016-2.27), the National Natural Science Foundation of China (NSFC #81271239, #81771205, #91632113), the Natural Science Foundation and Major Basic Research Program of Shanghai (16JC1420500, 16JC1420502), and the CAMS Innovation Fund for Medical Sciences

(CIFMS #2017-I2M-3-008). The funders had no role in study design, data collection and analysis, decision to publish, or preparation of the manuscript.

